# Pre-meiotic pairing of homologous chromosomes during *Drosophila* male meiosis

**DOI:** 10.1101/2021.12.07.471586

**Authors:** Thomas Rubin, Nicolas Macaisne, Ana Maria Vallés, Clara Guilleman, Isabelle Gaugué, Jean-René Huynh

## Abstract

In the early stages of meiosis, maternal and paternal chromosomes pair with their homologous partner and recombine to ensure exchange of genetic information and proper segregation. These events can vary drastically between species and between males and females of the same species. In *Drosophila,* in contrast to females, males do not form synaptonemal complexes (SCs), do not recombine and have no crossing-over; yet, males are able to segregate their chromosomes properly. Here, we investigated the early steps of homologues pairing in *Drosophila* males. We found that homologues are not paired in germline stem cells (GSCs) and become paired in the mitotic region before meiotic entry, similarly to females. Surprisingly, male germline cells express SC proteins, which localize to centromeres and promote pairing. We further found that the SUN/KASH (LINC) complex and microtubules are required for homologues pairing as in females. Chromosome movements are however much slower than in females and we demonstrate that this slow dynamic is compensated in males by having longer cell cycles. In agreement, slowing down cell cycles was sufficient to rescue pairing-defective mutants in female meiosis. Our results demonstrate that although meiosis differs significantly between males and females, sex-specific cell cycle kinetics are integrated with similar molecular mechanisms to achieve proper homologues pairing.

## INTRODUCTION

Meiosis is a two-step cell division process that generates haploid gametes from diploid parental germ cells (Bhalla & Dernburg, 2008; Zickler & Kleckner, 2015). During the first stages, homologous chromosomes face the daunting task of finding the correct homologue in the nuclear space, in order to recombine and exchange genetic information. This requires chromosomes to move, assess their homology, align along their length and pair. Pairing is then stabilized by the assembly of the synaptonemal complex (SC), which synapses both homologues together (Cahoon & Hawley, 2016). Double-strand breaks (DSBs) are formed on parental chromosomes and some of these DSBs are repaired with the homologous chromosome leading to the formation of crossing-overs (COs) and the reciprocal exchange of parental genetic information. The whole process is highly conserved in all sexually reproducing organisms from yeasts to humans. However, the underlying mechanisms allowing to achieve meiosis are remarkably diverse from one species to the next (Gerton & Hawley, 2005). For example, although chromosome movements to promote pairing is a common theme in many species, these movements are actin-dependent in *S. cerevisiae*, while they depend on microtubules in *S. pombe* (Rubin et al, 2020). Chromosomes move individually in *C. elegans*, whereas they follow global nuclei rotations in mouse and in *Drosophila*. Chromosomes are anchored to the nuclear membrane by centromeres in flies, but by telomeres in mice. Interestingly, this diversity of strategies between species also extends within the same species between male and female germ cells (Cahoon & Libuda, 2019). Indeed, beyond the obvious cellular dimorphism of a large female oocyte and small male sperm, there are many sex-specific differences in the meiotic process *per se*.

Chromosomal axis and SC are known to have different length in males and females of mice, worms and plants. Similarly, the number of COs and the response to DNA damage are sex-specific in many species (Cahoon & Libuda, 2019). *Drosophila* males are an interesting case, as in contrast to females, they do not form a synaptonemal complex nor make any crossing-over (McKee et al, 2012). However, males segregate their homologous chromosomes just fine. How males are able to associate their homologues without SC and recombination is not known.

In male flies, meiosis starts at the anterior region of the testis (Figure 1A). At the very anterior tip, Germline Stem Cells (GSCs) produce germ cells throughout the adult life (Fuller, 1993). GSCs divide asymmetrically to produce Gonioblasts (GBs), which go through four rounds of mitosis with incomplete cytokinesis to produce germline cysts made of 16 spermatogonia. All 16 cells remain connected by ring canals and a germline-specific structure called the fusome, which is made of ER-derived vesicles. The branched shape of the fusome allows to identify GSCs, GBs, and 2-, 4-, 8- and 16-cell cysts (cc) in the mitotic region. These mitoses are followed by an extended G2 phase marked by a 25-fold increase in cell size.

**Figure 1.**
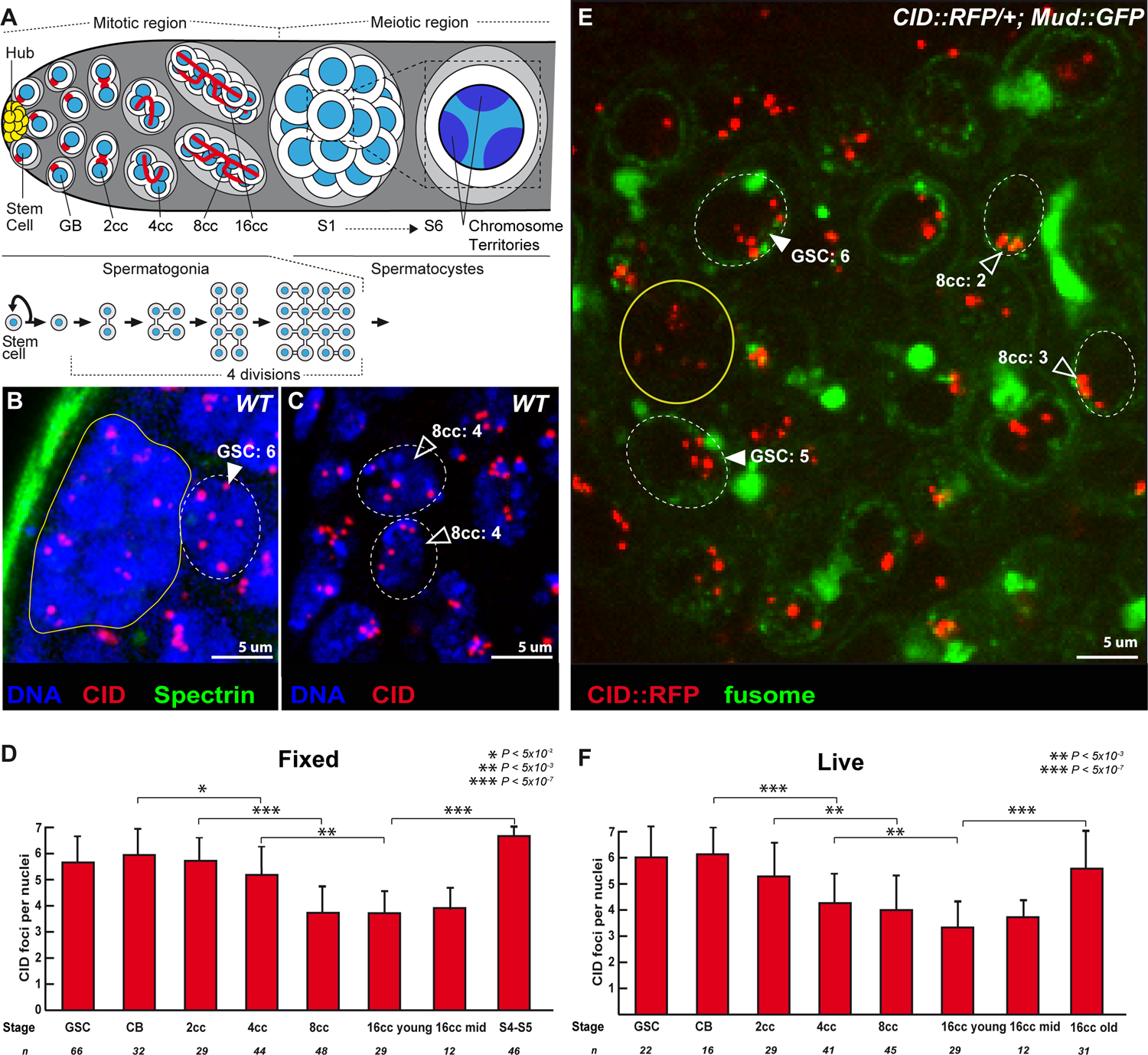
Unpaired centromeres in male germline stem cells become paired during cyst divisions. (A) Schematic overview of germ cell development in *Drosophila* testis. At the apical tip are the somatic hub cells (yellow), which serve as niche for germline stem cells (GSCs). GSCs divide asymmetrically giving rise to a GSC and a daughter cell, the gonioblast (GB), which undergoes four mitotic divisions by incomplete cytokinesis to produce interconnected 2-, 4-, 8-, and 16-cell spermatogonial cysts. Germ cells undergo an extended growth phase (S1-S6) entering meiosis as primary spermatocytes. A zoom view of chromosome territories in one spermatocyte. The spectrosome (red circles) of GSCs and GB, develops into a branched structure called fusome (red) during each division. (B,C) Z-section projections of wild-type fixed testes stained for centromeres (CID, red), fusome (α-Spectrin, green), and DNA (DAPI, blue). There are 6 centromeres in the GSC (B, arrowhead) but fewer in 8-cell cyst nuclei (C, open arrowheads), as quantified. (E) Z-section projection obtained by live-imaging of a testis expressing the centromere marker CID::RFP (red) and the fusome marker MUD::GFP (green). Two GSCs (arrowheads) identified by their position next to the hub cells (yellow circle) and an 8-cell cyst (8cc), with cells linked by the fusome. Two nuclei of an 8-cell cyst (8cc, open arrowheads), with fusome GFP, are marked by dotted lines. (D,F) Graphs plot the number of CID foci for each developmental stage in the mitotic region of fixed (E) or living (F) wild type testes. The number of analyzed cells is given below each stage. In B, C, D dotted lines surround germ cell nuclei. Scale bars in B, C, D are 5 µm. * p<5×10^-2^, ** p<5×10^-3^, *** p<5×10^-7^ (two-tailed Student’s t-test).

During this growth phase, homologous chromosomes are fully paired and then separate into chromosome territories (McKee et al, 2012). Pairing is then relaxed in these territories until chromosomes condense and form bivalents at metaphase I. Homologous chromosomes then segregate at anaphase I. At the spermatogonia-spermatocyte transition by the end of the mitotic region (Figure 1A), homologous chromosomes were shown to be highly paired (Vazquez et al, 2002). It is, however, not known when and how they become paired. Here, we investigated whether the paired state is passively inherited from embryonic stages or if there is *de novo* pairing in earlier stages of germ cell development in adults.

We and others have shown in *Drosophila* females that although chromosomes start meiosis already paired in the oocyte, female GSCs have unpaired chromosomes that become paired *de novo* during the four mitotic divisions (Christophorou et al, 2013; Joyce et al, 2013; Rubin et al, 2016). We further showed that homologues pairing requires extensive rotations of germ cells nuclei, and that these movements are driven by cytoplasmic microtubules (Christophorou et al, 2015). Microtubules forces are transmitted to the nuclear envelop and chromosomes by the SUN/KASH (LINC) complex encoded by the *klaroid* (*koi*) and *klarsicht* (*klar*) genes in *Drosophila*.

In this study, we investigated the pairing of homologous chromosomes in males before meiotic entry, i.e. in mitotic germ cells.

## RESULTS

### 1) Centromeres are unpaired in male germline stem cells and become progressively associated in the mitotic region

Germ cells were stained for α-Spectrin, which marks the fusome and allows identification of GSCs and the different stages of cysts differentiation (Figure 1A). To analyze chromosomes organization, we labeled centromeres with an antibody against CID, the *Drosophila* homologue of Cenp-A. *Drosophila* diploid cells have eight chromosomes forming four pairs of homologues. If all homologues are paired, one should count 4 foci of CID, whereas if some chromosomes are unpaired, there should be more than 4 (Takeo et al, 2011; Tanneti et al, 2011). We found in GSCs and GBs an average of 6 dots of CID (Figure 1B, 1D), which indicated that most chromosomes were unpaired at these stages. The number of CID dots went down to 4, in 8- and 16-cell cysts revealing that most chromosomes were associated at the end of the mitotic region (Figure 1C, 1D). In females, non-homologous centromeres can cluster with those of homologous pairs to form 1 or 2 dots of CID (Takeo et al, 2011; Tanneti et al, 2011). In contrast, we found that in males centromeres rarely clustered and remained in pairs (Figure 1D). Later during the growth phase in spermatocytes, when homologous chromosomes separate into different territories, the number of CID dots increased to 6, as previously shown (Figure 1D). We confirmed the results obtained on fixed testis by live-imaging using a CID-RFP transgene to track centromeres and a fusome marker tagged with GFP (Figure 1E, F, Movie S1-3).

We concluded that male centromeres are unpaired in GSCs and become *de novo* associated in pairs at the mitotic region similarly to females. However, they do not cluster.

### 2) Centromeres of chromosome II and III associate with their homologs in the mitotic region

Next, we analyzed whether pre-meiotic pairing at centromeres occurred between homologous chromosomes by labelling pericentromeric repeated sequences specific for each chromosome. We performed Fluorescent In Situ Hybridization (FISH) with the AACAC and dodeca probes, which identify pericentromeric regions of chromosome II and III, respectively (Figure 2). We considered that chromosomes were paired when only one focus was visible or when two foci were visible separated by a distance ≤0.70 um (Blumenstiel et al, 2008; Gong et al, 2005).

**Figure 2.**
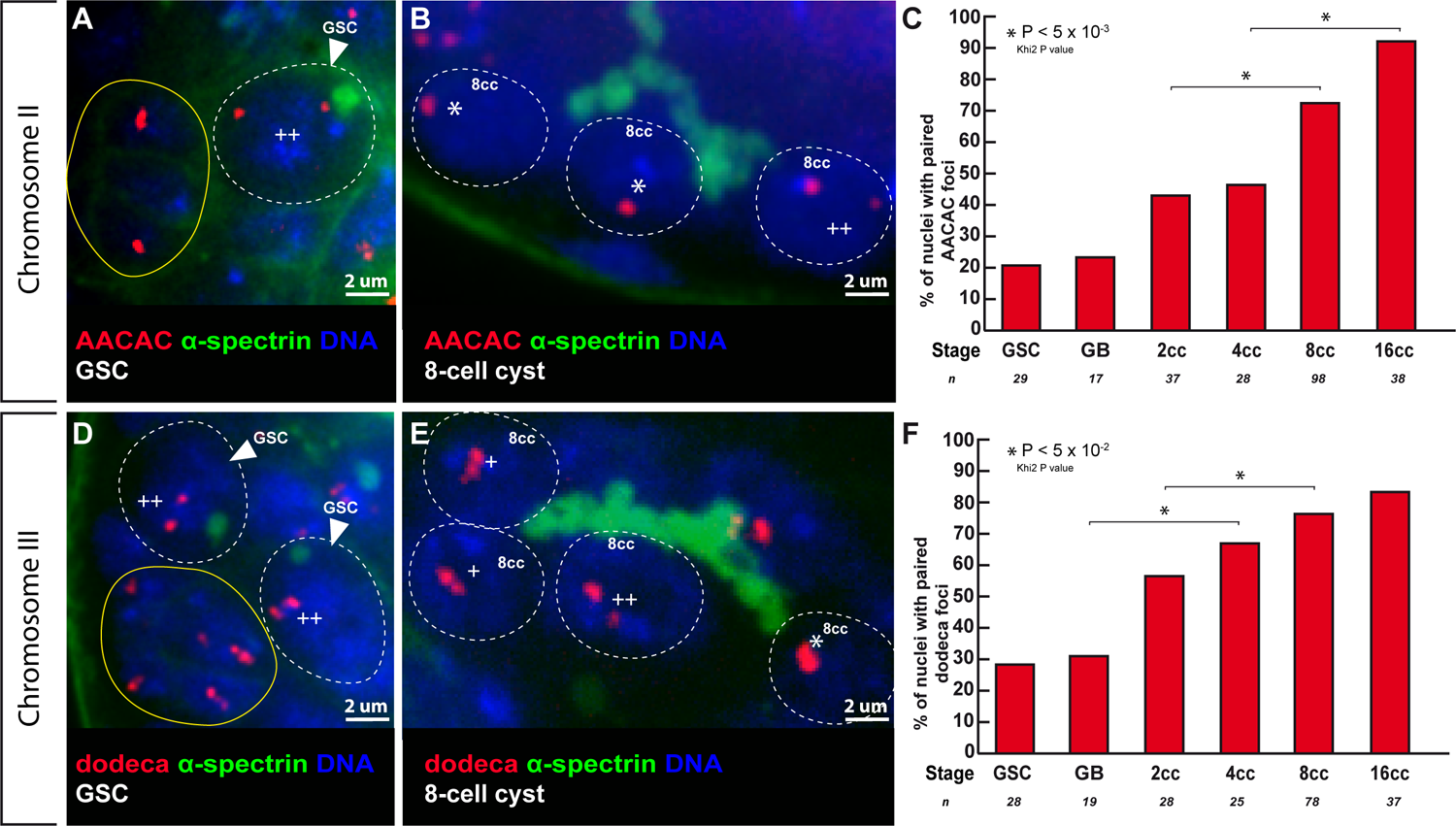
Centromeres of chromosome II and III become associated in the mitotic region. (A, B) Confocal Z-section projections of wild-type testes stained for chromosome II centromeres probe (AACAC, red), fusome (α–spectrin, green) and DNA (DAPI, blue). (A) GSC shows 2 foci separated by a distance > 0,7 µm (++), thus unpaired; hub is marked in yellow. (B) 8-cell cyst shows 2 nuclei with one focus (*), thus centromeres are paired and 1 nucleus with two foci (++) separated by a distance > 0,7 µm and thus unpaired. (C) Graphs plot the percentage of paired chromosome II centromeres for each developmental stage in the mitotic region using the AACA probe. The number of cells analyzed is indicated under each stage.* p<0.05 (khi2 test comparing 2cc with 8cc and 4cc with 16cc). (D, E) Confocal Z-section projections of wild-type testes stained for chromosome III centromere probe (dodeca, red), fusome (α–spectrin, green) and DNA (DAPI, blue). (D) 2 GSCs with 2 foci separated by > 0,7 mm (++), thus unpaired; hub is marked in yellow. (E) 8-cell cyst shows a nucleus with one foci (*), thus paired, 2 nuclei with two foci (+) separated by a distance of ≤ 0,7 µm, thus paired, and 1 nucleus with 2 foci (++) separated by a distance of 0,7 µm, thus unpaired. (F) Graph plots percentage of paired chromosome III centromeres for each developmental stage in the mitotic region using the dodeca probe. The number of cells analyzed is indicated below each stage. In A, B, D, E dotted lines surround germ cell nuclei. Scale bars are 2 µm. * p<0.005 (khi2 test comparing GB with 4cc and 2cc with 8cc).

With both probes, we found that less than 30% of chromosomes II and III were paired in GSCs, that went up to 90% (n=38) and 80% (n=37) for chromosome II and III respectively in 16-cell cysts. We concluded that centromeres of chromosomes II and III become associated with their homologues in the mitotic region.

Sex chromosomes X and Y share little homologous sequences outside the clusters of rDNA genes. The intergenic spacer (IGS) region located upstream of each rDNA repeats was shown to be necessary and sufficient for X and Y pairing (Mckee & Karpen, 1990). We used a probe labelling the IGS to investigate sex chromosomes behavior. We found that it marked one unique but large sub-region of the nucleus from GSCs to 16-cell cysts (Fig S1). This region was too broad to allow us to conclude on X/Y pairing (McKee et al, 2012). However, as in females, we counted between 6 and 7 dots of centromeres in GSCs and not 8 (Figure 1D), indicating that some chromosomes were paired even in GSCs. In females, X-chromosomes are already paired in GSCs (Christophorou et al, 2013; Joyce et al, 2013); it is thus possible, although it remains to be demonstrated, that in males the X and Y-chromosomes are also paired in GSCs and cyst cells.

### 3) Synaptonemal complex components are expressed in male germ cells, localize at centromeres and chromosome arms, and are required for efficient pairing of homologues

In *Drosophila* females, we showed that pre-meiotic pairing of autosome centromeres depend on components of the SC such as C(3)G and Corona (Cona), which localized at centromeres in the mitotic region (Christophorou et al, 2013). Interestingly, although no SC structure has ever been described in males, both the FlyAtlas2 and modENCODE datasets detected *cona*, and to a lesser extent *c(3)G*, expression in testis. We confirmed the expression of *c(3)G* in testis by RT-qPCR (Figure S2A). We used an antibody against C(3)G to determine the localization of the endogenous protein, and found that C(3)G associated to chromosomes in the mitotic region (Figure 3A). All signals disappeared completely in *c(3)G* mutant germ cells, demonstrating the antibody specificity (Figure 3B). Super-resolution microscopy revealed that C(3)G formed small foci along chromosome arms (Figure 3E). The largest and brightest foci were associated with centromeres as in female germ cells (Figure 3C, D and S2D-G). We validated these findings with a tagged C(3)G-GFP that colocalized with Cid-RFP by live-imaging (Figure S2B-C, Movie S6).

**Figure 3.**
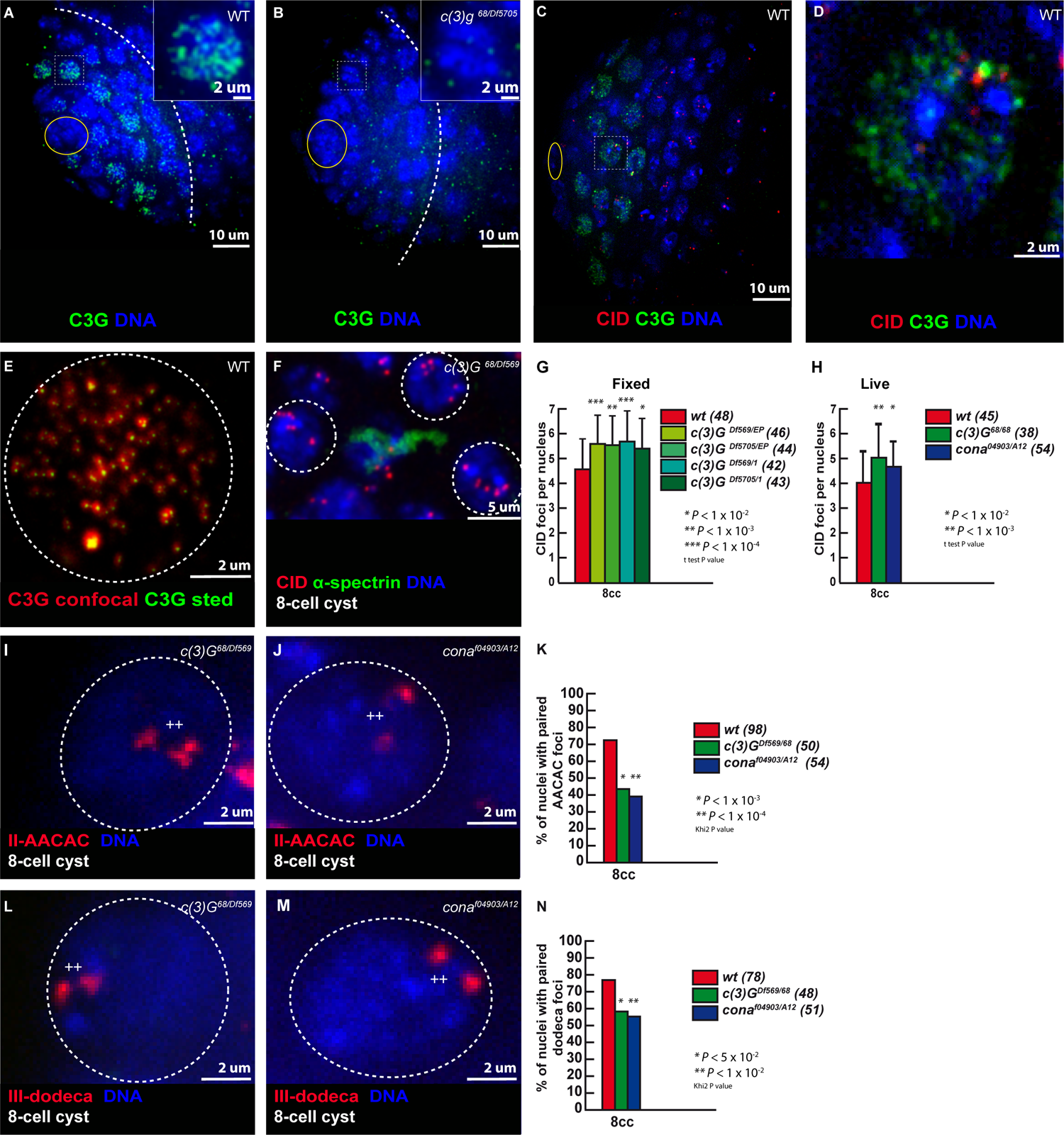
The Synaptonemal Complex component C(3)G is present in the mitotic region and promotes centromere association. (A, B) Confocal Z-section projections of wild-type and *c(3)G^68^*/*Df5705* (B) testes stained for C(3)G (green) and (DAPI, blue). Dotted white lines delimit the mitotic region; hub is marked by the yellow circle. (C,D) Confocal Z-section projections of wild-type testes stained for centromeres (CID, red), C(3)G (green) and DNA (blue); hub is marked by a yellow circle. Inserts in A, B show magnified nuclei corresponding to outlined nuclei. Outlined nucleus is magnified in (D). Scale bars in A, B are 10 µm and inserts are 2 µm. (E) Z-section projection acquired using STED microscopy of a WT 2-cell cyst stained for C(3)G (red is confocal, green is STED). Scale bar is 2 µm. (F) Confocal Z-section projection of an 8-cell cyst from *c(3)G^68^*/*Df(3R)BSC569* stained for centromere (CID, red), fusome (α-Spectrin, green), and DNA (DAPI, blue). Three nuclei (dotted circles) show unpaired centrosomes. Scale bar is 5 um. (G,H) Graphs plot the number of CID foci of 8-cell cysts in fixed wild-type, *Df(3R)BSC569*/*c(3)G^EP^*, *Df5705/c(3)G^EP^*, *Df(3R)BSC569*/*c(3)G^1^*, *Df5705/c(3)G^1^* (G) and in live wild-type, *c(3)G^68^/c(3)G^68^* and *cona^A12^/cona^04973^* (H) testes. The number of analyzed cells is indicated next to each genotype. * p<0.01, ** p<0.001, *** p<0.0001, (two-tailed Student’s t-test). (I, J) Confocal Z-section projections of *c(3)G^68^*/*Df(3R)BSC569* (I) and *cona^A12^/cona^04973^* (J) 8-cell cyst nuclei stained for dodeca (red) and DNA. In both cases, nuclei (dotted circles) show two foci separated by > 0.7 mm (++) indicating unpaired centromeres. (K) Graph plots the percentage of paired chromosomes III in 8-cell cysts using the dodeca probe in wild-type, *c(3)G^68^*/*Df(3R)BSC569* and *cona^A12^/cona^04973^* fixed testis. The number of analyzed cells is indicated next to each genotype. * p<0.05, ** p<0.01 (khi2 test). (L, M) Confocal Z-section projections of *c(3)G^68^*/*Df(3R)BSC569* (L) and *cona^A12^/cona^04973^*. (M) 8-cell cyst nuclei stained for AACAC (red) and DNA. Both nuclei (dotted circles) show two foci separated by > 0.7 µm (++) indicating unpaired centromeres. Scale bars in I, J, L, M are 2 µm. (N) Graph plots the percentage of paired chromosomes II in 8-cell cysts using the AACAC probe in wild-type, *c(3)G^68^*/*Df(3R)BSC569* and *cona^A12^/cona^04973^* fixed testes. The number of analyzed cells (n) is indicated next to each genotype. * p<0.001, ** p<0.0001 (khi2 test). Chromosome II and III centromeres are considered paired when the distance between 2 foci ≤ 0.7 µm.

To investigate *c(3)G* and *cona* function in male chromosome organization, we counted the number of CID foci in four different *c(3)G* mutant backgrounds at the 8-cells cyst stage. We found an average of 5,5 dots of CID, which was significantly higher than in wild type germ cells (Figure 3G). We confirmed this difference by live-imaging in *c(3)G* and in *cona* mutant cells (Figure 3H). We next assessed the number of paired centromeres of chromosome II and III by FISH. In *c(3)G* and *cona* mutant cells, less than 40% of chromosome II were paired and less than 50% for chromosome III compared to more than 70% in wild type 8-cell cysts (Figure 3I-N).

We concluded that SC components are expressed in the male mitotic region and are necessary for efficient pairing of homologous centromeres.

### 4) Dynamic movements of centromeres depend on microtubules and Dynein motor

A common theme in meiosis is to set chromosomes in motion so that homologues have a chance to meet, assess their homology and pair. In *Drosophila* females, we showed that extensive rotations of germ cells nuclei increased the efficiency of chromosome pairing (Christophorou et al, 2015). In males, we followed chromosomes movements using a CID-RFP transgene and calculated the relative covered volume per second for each centromere by live-imaging as previously done (Figure 4A, B) (Christophorou et al, 2015). We found that centromeres were dynamic and followed directed motions within the nuclear space. These movements were more prominent at the 4- and 8-cell stages that coincided with the increase in chromosome pairing that we detected (Figure 4C-F). However, even at the 8-cell stage, the relative volumes covered by centromeres were markedly smaller than in females (Figure 4G). For example, we never detected complete nuclear rotations of centromeres during the periods of recording. On average, we found that at the 8-cell stage the nuclear coverage in males was 3 times smaller than in females (Figure 4G).

**Figure 4.**
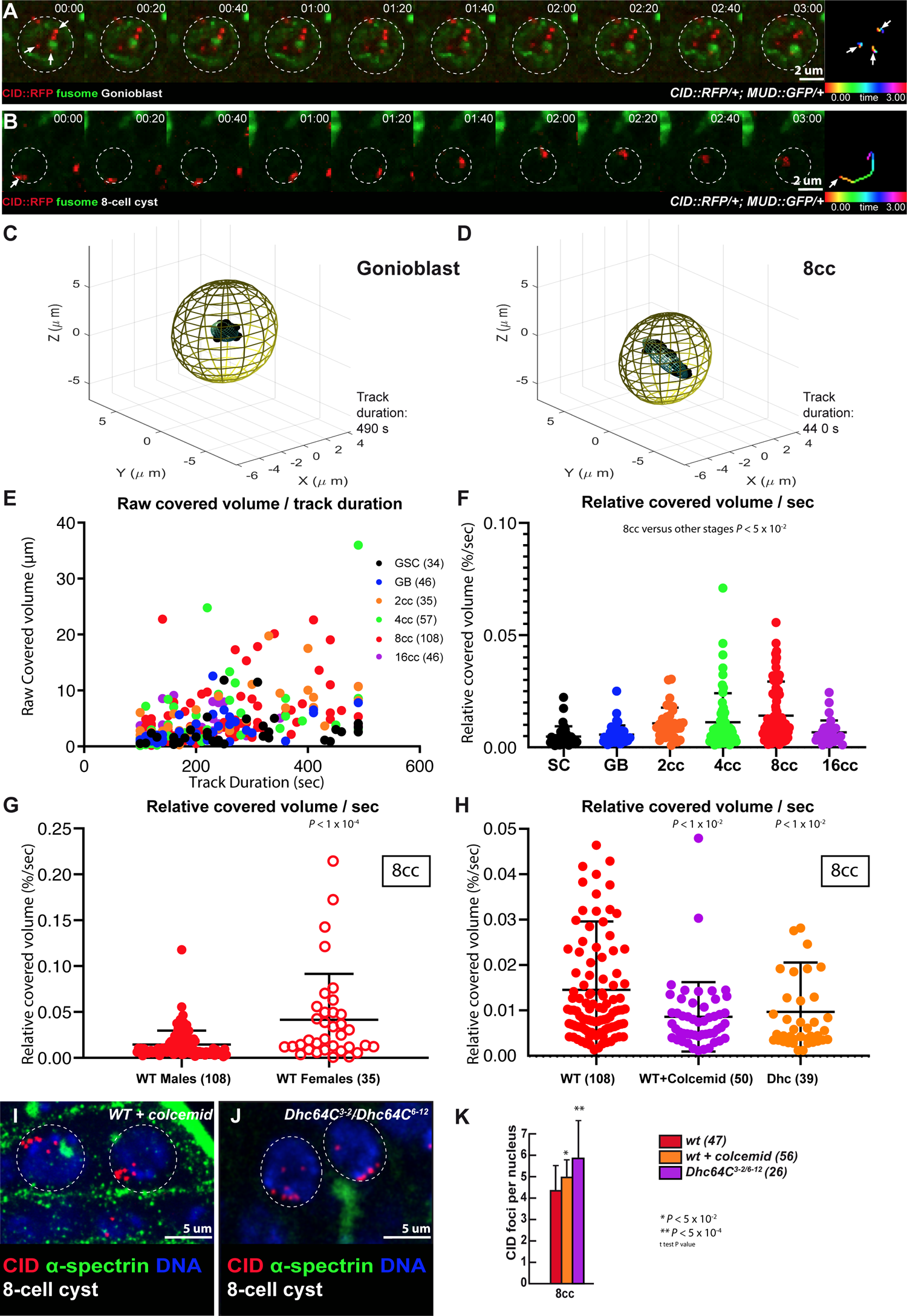
Centromere pairing of 8-cell cysts exhibit dynamic rotations mediated by microtubules and dynein. (A, B) Selected projections of nuclei from a GB (A) and an 8-cell cyst (B) over a 3-min time course (nuclear areas are indicated by dotted circles). Time-colored tracking of three CID–RFP dots (arrows) in (A) and one CID–RFP dot (arrowhead) in (B) are shown in the rightmost panels. (C, D) Three-dimensional representations of the covered volume of centromeres in time from one representative track; an ellipsoid was arbitrarily centered representing the nuclear volume in a GB (C) or an 8-cell cyst (D). (E) Raw covered volumes of each centromere track from GSC, GB, 2-to 16-cell cyst as a function of track duration. The number of tracks analyzed is given next to each cell type. (F) Relative covered volume (raw covered volume/nuclear volume) per second for each track at different cyst stages (mean ± s.d., Mann–Whitney U-test comparing the wild-type 8-cell cyst with other stages P <0.05). (G) The relative covered volume (raw covered volume/nuclear volume) per second in 8-cell cyst nuclei of males is much less when compared to females. (mean ± s.d., Mann–Whitney U-test, P <0.0001; The number of foci analyzed is indicated. (H) The relative covered volume (raw covered volume/nuclear volume) per second in 8-cell cyst nuclei of males treated with colcemid or in *Dhc* mutant is strongly reduced as compared to wild-type. (mean ± s.d., Mann–Whitney U-test, P <0.01; WT: n = 108 centromeric foci; WT + colcemid: n=50 centromeric foci; *Dhc*: n=39 centromeric foci). (I, J) Confocal Z-section projections of testes from wild-type flies treated with colcemid (wt+colcemid) (I) and *Dhc^3-2^/Dhc^6-12^* (J) stained for centromeres (CID, red), fusome (α–spectrin, green) and DNA (DAPI, blue). 8-cell cyst nuclei are indicated by dotted circles. Note that in both conditions centromeres are unpaired. Scale bars are 5 µm. (K) Graph plots the number of CID foci in 8-cell cyst nuclei in wild-type (wt), wt + colcemid and in *Dhc^3-2^/Dhc^6-12^* mutant. The number of analyzed cells is indicated next to each condition (two-tailed Student’s t-test, * p<0.05, ** p<0.0005.)

We next tested whether these movements depended on the microtubule cytoskeleton and its associated motors. We fed male flies with the microtubule-polymerization inhibitor colcemid for 4h and then dissected the testis for live-imaging. We found that centromere movements were markedly reduced in the presence of colcemid (Figure 4H, Movie S5). Likewise, in dynein *Dhc64C* mutant males, centromere dynamics was strongly reduced (Figure 4H, Movie S4). Importantly, in both cases, centromere pairing was strongly affected at the 8-cell stage (Figure 4I-K).

We concluded that centromeres are dynamic in males, albeit much less than in females. Nevertheless, as in females, these movements depend on microtubules and are required for efficient pairing of centromeres.

### 5) Drosophila LINC complex is required for efficient centromere pairing in males

The LINC complex is formed by two transmembrane proteins, which bridge the inner and outer nuclear membranes with the cytoplasmic cytoskeleton. In *Drosophila*, there are two SUN-domain proteins, Klaroid (Koi) and Spag4, localizing at the inner nuclear membrane; and two KASH-domain proteins, Klarsicht (Klar) and MSP-300, localizing at the outer membrane. We generated GFP knock-in lines by CRISPR at the endogenous *klaroid* and *klarsicht* loci to investigate their localization. We also used an independent GFP-trap line inserted into the *klaroid* locus. All three lines showed a homogeneous localization at the nuclear envelop (NE) in GSCs, GBs and 2-cell cysts (Figure 5A, B, E, F). However, both Klar and Koi also formed 1 or 2 dots at the NE of 4-, 8- and early 16-cell cysts (Figure 5A, B, E, F, arrowheads). Localization of Koi and Klar then became homogenous again at the NE in later stages. Interestingly, we noticed that Koi and Klar dots were often associated with centromeres at the NE (Figure 5C, D, G, H and Figure 5I-L). This co-localization of Klar, Koi and centromeres suggested that the LINC complex could be involved in centromere pairing in males.

**Figure 5.**
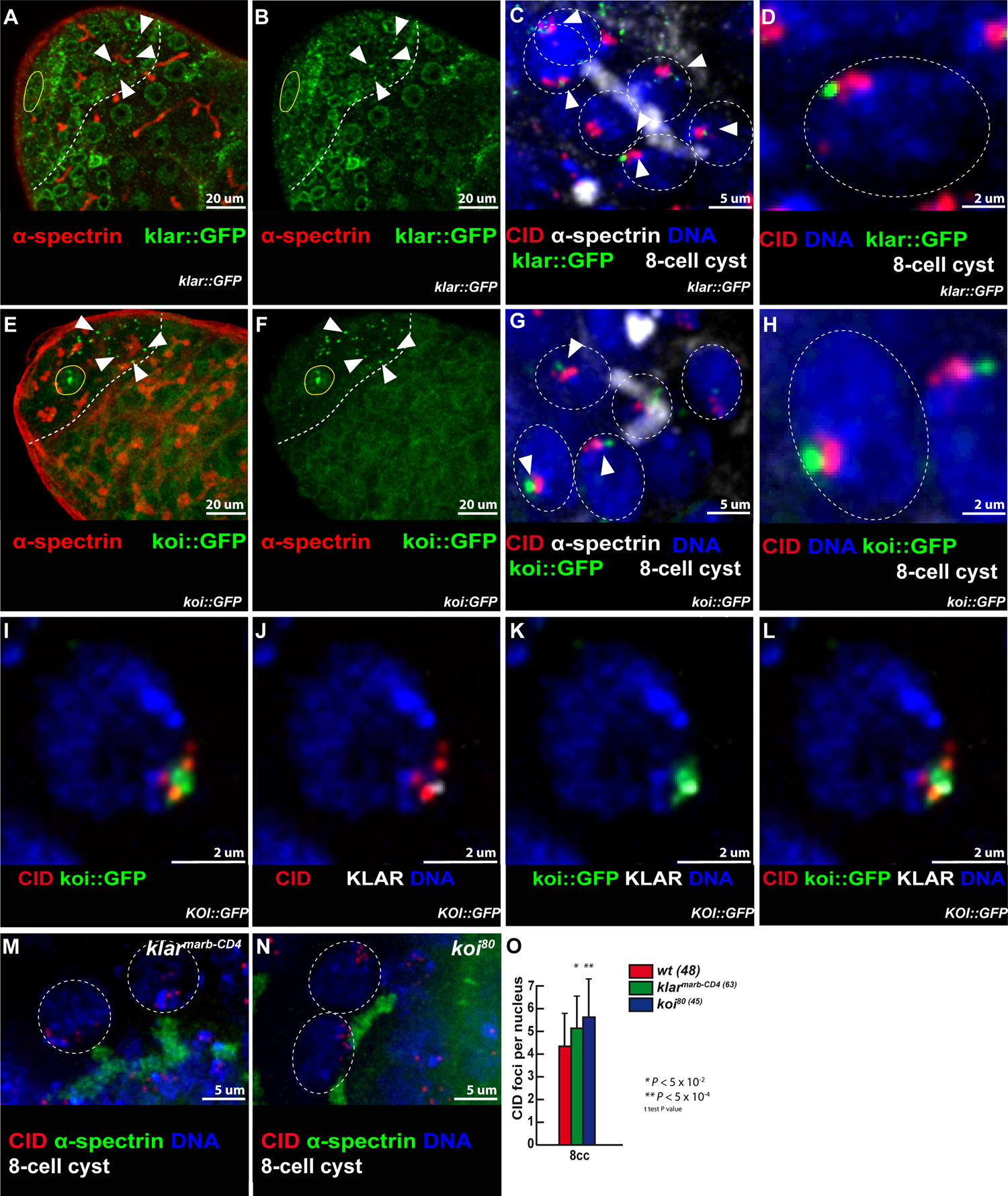
Klarsicht and Klaroid localize near centromeres in the mitotic region and are required for 8-cell cyst chromosome pairing in males. **(A, B) Confocal Z-section** projections of klar::GFP (green) stained for fusome (α-Spectrin, red). In A, B, E, F dotted lines delimit the mitotic region and the hub is marked by the yellow circle. Arrowheads point to Klar::GFP dots at the NE. Scale bars are 20 µm. (C) Confocal Z-sections projection of Klar::GFP (green) stained for fusome (α-Spectrin, white), centromeres (CID, red) and DNA (DAPI, blue). (D) Magnified view of a nucleus outlined in (C) showing CID–Klar::GFP association. (E, F) Confocal Z-section projections of koi::GFP (green) stained for fusome (α– spectrin, red). Arrowheads point to Klar::GFP dots at the NE. (G) Confocal Z-sections projections of koi::GFP (green) stained for fusome (α–spectrin, white), centromeres (CID, red) and DNA (DAPI, blue). (H) Magnified view of a nucleus outlined in (G) showing CID– Koi::GFP association. (I-L) Confocal Z-section projections of a nucleus from an 8-cell cyst stained for centromere (CID, red), Klarsicht (Klar, grey), Klaroid::GFP (Koi, green) and DNA (DAPI, blue); scale bars 2um. (M, N) Confocal Z-section projections of *klar^marb-CD4^*/*klar^marb-CD4^* (M) and *koi^80^/koi^80^* (N) stained for centromeres (CID, red), fusome (α–spectrin, green) and DNA (DAPI, blue). Scale bars are 5 µm. In C, D, G, H, M, N, 8-cell cyst nuclei are indicated by a dotted circle in each image. Scale bars are 5 µm. (O) Graph plots the number of CID foci in 8-cell cyst nuclei from wild-type, *klar^marb-CD4^*/*klar^marb-CD4^* and *koi^80^/koi^80^* mutants. The number of analyzed cells is indicated next to each genotype (two-tailed Student’s t-test, * p<0.05, ** p<0.0005.)

We tested a potential function of SUN/KASH proteins by analyzing centromere pairing in the mutants’ background at the 8-cell stage. We found an average of 5,7 dots of CID in *koi* mutants, and 5 dots in *klar* mutant germ cells, in contrast to 4 in wild type control flies (Figure 5M-O). We did not investigate Spag4, as *spag4* is not expressed in early male germ cells (Shi et al, 2020) and we did not test *MSP-300* mutants, as MSP-300 is known to interact with the actin cytoskeleton (Volk, 1992; Yu et al, 2006).

These results showed that the *Drosophila* LINC complex (Klar/Koi) is required for efficient pairing of chromosomes in males.

### 6) Mitotic cell cycle durations in males are much longer than in females

Our results revealed that chromosome pairing in males is pre-meiotic, depends on SC components and on movements driven by the microtubule cytoskeleton and associated proteins, which is strikingly similar to females. Yet, centromere dynamics are much slower in males than in females. For comparison, the average covered volume per second for centromeres in wild type males is less than in *klaroid* mutant females, which showed clear pairing and synapsis defects (0,014%/sec in wild type males vs 0,029%/sec in koi females) (Christophorou et al, 2015). Thus, how could slow movements be sufficient for pairing in males but not in females, despite both using identical molecular machinery?

A first clue to this paradox came when we measured the relative volumes covered by centromeres in males. We noticed that the nuclear volumes of 2-, 4-, 8- and 16-cell cysts were larger in males than in females (Figure S3A). It suggested to us that cells had more time to grow in males than in females during each interphase. To test this hypothesis, we determined the length of time spent at each stage of germ cell differentiation in wild type males and females (See Experimental Procedures). We measured the frequency of mitosis at each stage by live-imaging using a GFP marker for mitosis (survivin-GFP). We recorded more than 254 hours of 18 testis, and 235 hours of 26 ovaries. We found that both female and male GSCs had a similar cell cycle duration of around 20h, as previously published (Morris & Spradling, 2011; Sheng & Matunis, 2011). However, in female cysts, cell cycles became shorter at each division, reaching 6h in 8-cell cysts (Figure 6A). In contrast, male cysts cell cycles were longer than in GSCs, with a maximum of 40h for 2-cell cysts (Figure 6A). At the 8-cell cyst stage, we measured 30h duration in males, which is about 5 times longer than in females. By adding all the durations from GSCs to 8-cell cysts, we found that the length of time for a GSC to reach the 8-cell stage lasted 48h in females and 148h in males on average (Figure 6B). The developmental window during which homologue chromosomes pair is thus about 3 times longer in males than in females. These observations support our hypothesis that slower chromosome movements in males could be compensated by an increase in the pairing period.

**Figure 6.**
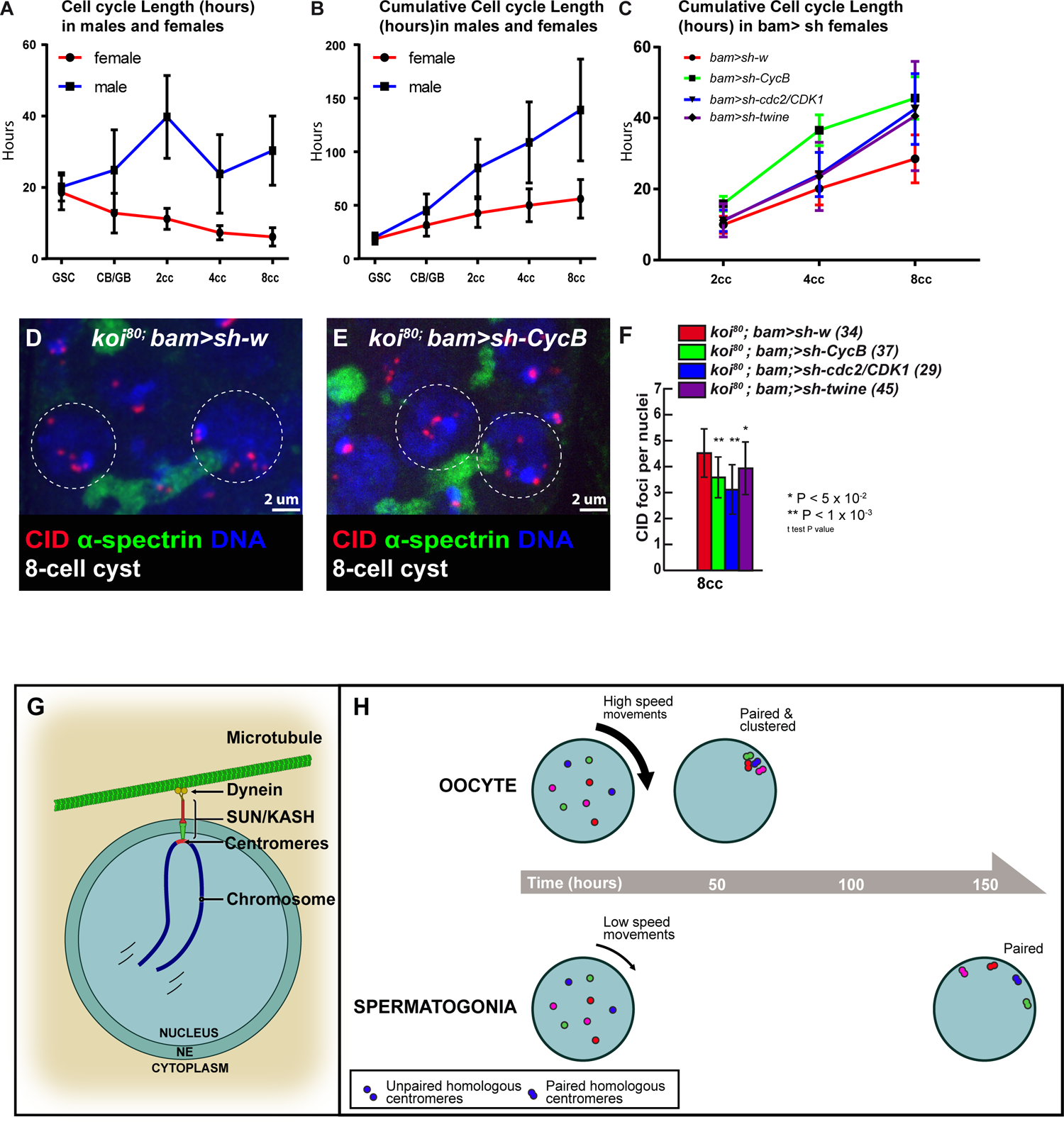
Slowing down the cell cycle rescues koi pairing defects in females. (A, B) Cell cycle length in males and females of developing cysts in the mitotic region represented as the number of hours spent per stage (A) or as the number of hours elapsed since GSC at each stage (B). (C) Cumulative cell cycle length hours in the mitotic region of females expressing a control *bam>sh-w (red)*, *bam>sh-CycB (green)*, *bam>sh-cdc2/Cdk1 (blue),* and *bam>sh-twine (black).* (D, E) Confocal Z-section projections of control *koi^80^/koi^80^; bam/sh-w* (D) and *koi^80^/koi^80^; bam/sh-CycB* (E) ovaries stained for centromeres (CID, red), fusome (α–spectrin, green) and DNA (DAPI, blue). 8-cell cyst nuclei are indicated by a dotted circle in each image. Scale bars are 2 µm. (F) Graph plots the number of CID foci in 8-cell cyst nuclei in *koi^80^/koi^80^; bam>sh-w koi^80^/koi^80^; bam>sh-CycB, koi^80^/koi^80^; bam>sh-cdc2/CDK1* and *koi^80^/koi^80^; bam>sh-twine*. The number of analyzed cells is indicated next to each genotype (two-tailed Student’s t-test, * p<0.05, ** p<0.001). (G) and (H) Graphical abstract.

### 7) Increasing cell cycle length is sufficient to rescue pairing defects in females

We wanted to test functionally the correlation between speed of chromosome pairing and cell cycle duration. We chose *klaroid* mutant females because centromeres are not attached to the NE, chromosome movements are inefficient and homologues pairing is defective (Christophorou et al, 2015). To test if increasing the cell cycle length would be sufficient to rescue pairing defects in *koi* females, we reduced the amount of Cyclin B, the limiting factor of the Cdk1/CycB complex, which promotes mitotic entry (Lindqvist et al, 2009). We targeted *cycB* only during the pairing window of 2- to 8-cell cysts by using the *bam* promoter to express an shRNA targeting *cycB* (*bam*>sh-cycB). In this partial knock-down (KD) condition, 16-cell cysts could still form normally. Indeed, after recording 37 germarium for 257 hours, we found that the duration of the pairing window had increased to 45h in *bam*>sh-cycB females compared to 28h in *bam*>sh-white control flies (Figure 6C). These mutant conditions thus allowed us to almost double the length of the pairing period.

Next, we counted the number of CID dots in double mutant *koi*, *bam*>sh-cycB females, compared to *koi*, *bam*>sh-white control flies. In 8-cell cysts, the number of CID foci was significantly reduced to 3,6 in *koi*, *bam*>sh-cycB females, compared to 4,5 in *koi*, *bam*>sh-white control flies (Figure 6D-F).

To confirm that lengthening the cell cycle was sufficient to rescue pairing defects, we tested two additional cell cycle regulators to increase cell cycle duration: either knocking down the key kinase Cdk1/Cdc2 (*bam*>sh-*cdc2*) or by knocking down twine/Cdc25 (*bam*>sh-*twe*), a phosphatase, which activates CycB/Cdk1 activity (Lindqvist et al, 2009). In *cdc2* KD, we found that the duration of the pairing window increased to 42h, and in *twe* KD to 41 hr, which was about a 50% increase of the wild type duration (Figure 6C). We found a strong reduction in the number of CID foci in *koi*, *bam*>sh-*cdc2* and a mild reduction in *koi*, *bam*>sh-*twe* females compared to *koi*, *bam*>sh-white control flies (Figure 6F).

We concluded that increasing cell cycle length in three independent genetic backgrounds was sufficient to partially rescue pairing defects in *klaroid* mutant females.

## DISCUSSION

*Drosophila* males have always been considered outliers for meiosis as they do not form synaptonemal complexes nor do they need recombination between chromosomes to segregate homologues. Here, our results demonstrate that they share many unsuspected similarities with females. Firstly, timing of pairing is identical, there is *de novo* homologue pairing in the mitotic region from GSCs to 16-cell cysts. Secondly, underlying molecular mechanisms are similar, as males also depend on the microtubule cytoskeleton, SUN/KASH proteins and SC components for efficient pairing.

There are, however, interesting differences. In females, the early localization of SC components at centromeres may initiate the formation of the SC along chromosome arms and the complete synapsis of homologues (Tanneti et al, 2011). Obviously in males, C(3)G and Corona at centromeres do not initiate the formation of the SC, and once homologues are paired, they are separated into different nuclear territories in spermatocytes. In these territories, homologues then become unpaired and centromeres dissociate. These are two different strategies to segregate homologues at metaphase I: in females, homologues are fully synapsed in the same nuclear space; whereas in males, homologues are unpaired but separated into different nuclear territories. Another interesting difference revealed by our study is the slower dynamics of male meiotic chromosomes as compared to females. Within the limitations of our live-imaging protocol, we never observed dramatic rotations of nuclei as we have described in females. Nevertheless, male centromeres follow the global movements of each nucleus. It remains thus possible that slow nuclear rotations occur, which could not be recorded within the timeframe of our experiments. These coordinated movements are thus similar to what we described in *Drosophila* females and to chromosome movements in mouse spermatocytes (Christophorou et al, 2015; Lee et al, 2015; Shibuya et al, 2014). In contrast, in *C. elegans*, individual chromosomes move independently, although they also rely on the LINC complex and microtubules (Hiraoka & Dernburg, 2009; Zetka et al, 2020).

Although, C(3)G and Corona do not initiate the formation of an SC in males, we found by super-resolution microscopy that they form many foci associated with centromeres and along chromosome arms, specifically in the mitotic region. It is tempting to speculate that these foci could be sites of pairing between homologues, and that they could act as “button loci” as in somatic cells (Viets et al, 2019). Indeed, homologue chromosomes are also paired in *Drosophila* somatic cells (Joyce et al, 2016). Recent studies have identified genomic regions, associated with TADs (Topologically Associated Domains), which mediate somatic pairing of homologues. These button loci were identified by DNA FISH, Hi-C and by biophysical modelling (Child et al, 2021; Rowley et al, 2019; Viets et al, 2019). Button loci are enriched with insulator or architectural proteins such as CTCF, Cp190 or Mod(mdg4) (Rowley et al, 2019). It would be interesting to test whether C(3)G and Corona associate with these proteins at button loci in germline cells. However, it remains a challenge to isolate enough pre-meiotic germline cells to perform co-IP or ChIP-seq experiments.

One major difference between male and female uncovered by our study is the difference in cell cycle durations. Previous studies had already reported that cell cycle regulation differs between males and females germ cells (Gadre et al, 2020; Hinnant et al, 2017; Insco et al, 2009). However, by analyzing fixed samples, these studies could only estimate the relative ratios of cell cycle phases between GSCs and the different cyst stages. Here, using a live-imaging approach similar to Sheng and Matunis (Sheng & Matunis, 2011), we obtained more direct measurements of cell cycle length, not only of GSCs but also of all cyst stages. Our results are in very good agreement with those obtained with fixed samples. For example, we found a peak of duration at the 2-cell stage in males, which lasted around 40hr, which is about twice as long as the 4-cell stage that we measured at 20hr. This is very similar to the 2:1 ratio found with fixed samples (Gadre et al, 2020). Overall, we found that the window when homologous chromosomes pair in males is 3 times longer than in females. These differences in timing and kinetics for the same process in males and females could well be widespread among species. For example, in *C. elegans*, prophase I of meiosis is completed in about 20h in males, while in oogenesis it lasts around 60hr (Cahoon & Libuda, 2019). This sexual dimorphism could explain different sensitivity in males and females to the same mutation affecting meiosis. Indeed, mutations of checkpoint proteins or the presence of unrepaired DNA breaks could have stronger consequences in germ cells having a faster cell cycle, males or females depending on the species, than in slower cells, which may compensate by having longer cell cycles. It is also possible that differences in cell cycle timing may impose different strategies for homologue pairing or synapsis between species.

For example, yeast *S. cerevisiae* has to pair 16 pairs of chromosomes in a few hours and requires DSBs, compared to *Drosophila* males, which have only 4 pairs of chromosomes and about 100 hours to achieve pairing and do not require DSBs. Our results show that time is an important dimension to understand the evolution of different meiotic strategies between sexes and species.

## ACKNOWLEDGMENTS

We would like to thank Scott Hawley for the initial discussion on the expression of synaptonemal complex genes in males and Isabelle Bonnet for her help with centromere tracking. We are grateful to M.T. Fuller, S. Hawley, S. Roth, Y. Bellaiche, Bloomington Stock Center, DSHB Hybridoma Center, for antibodies and flies. We are grateful to the Orion Imaging facility at CIRB (Collège de France). JRH lab is supported by CNRS, Inserm, Collège de France, FRM (Equipe FRM DEQ20160334884), ANR (ANR-13-BSV2-0007-02 PlasTiSiPi; ANR-15-CE13-0001-01, AbsCyStem) and the Bettencourt-Schueller foundation (FSER).

## Supplementary Figures

**Supplementary Figure 1.**
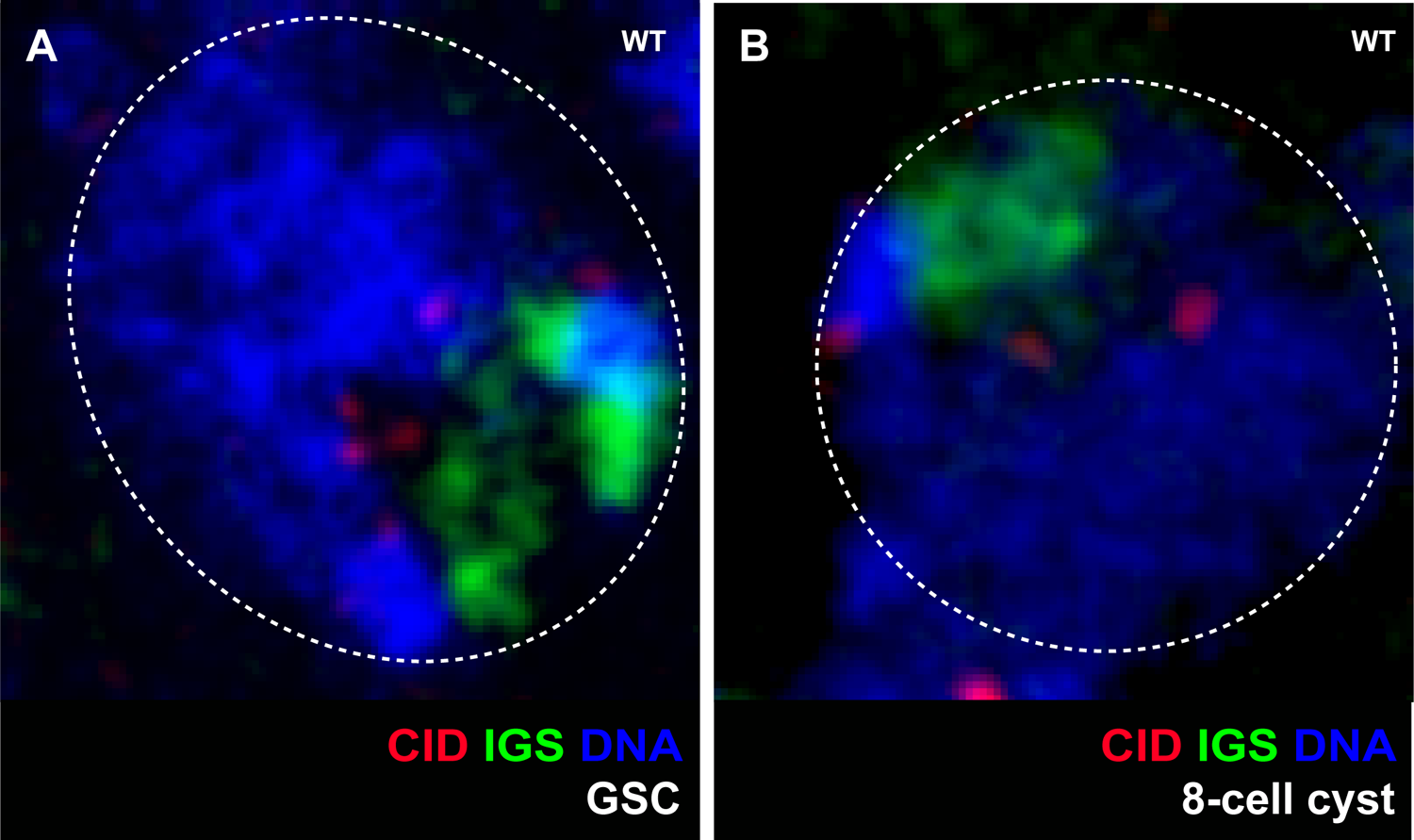
(A, B) Confocal Z-section projections of a wild-type testis stained for chromosome rDNA (IGS probe, green), centromere (CID, red) and DNA (DAPI, blue). (A) GSC shows a broad and diffuse IGS signal next to CID foci. (B) 8-cell cyst shows broad and diffuse IGS signal next to CID foci. Dotted lines surround nuclei.

**Supplementary Figure 2.**
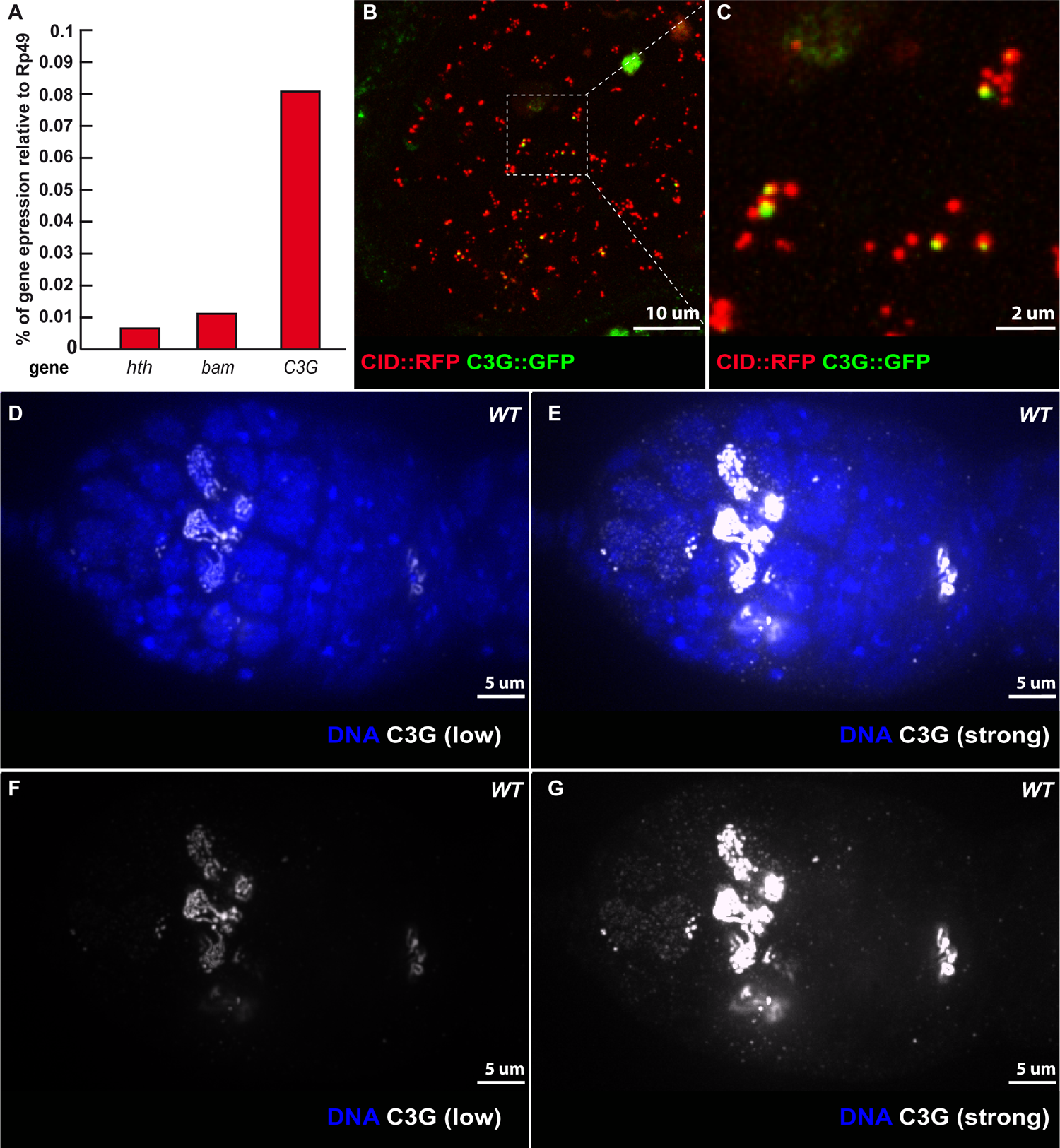
(A) Real-time PCR of *hth*, *bam* and *c(3)G.* Relative amount of gene expression with respect to *rp49* in testis. (B) Z-section projections obtained by live-imaging of a testis expressing CID::RFP (red) and c(3)G::GFP (green). (C) Magnified view of a CID::RFP–C(3)G::GFP association corresponding to region outlined in (B). (D-G) Confocal Z-section projections of a wild-type germarium stained for C(3)G (white) and DNA (DAPI, blue). Low signal panels (D, F) show the classical rod pattern of synaptonemal complex in meiotic region, and High signal panels (E,G) reveals C(3)G nuclear localization in the mitotic region.

**Supplementary Figure 3.**
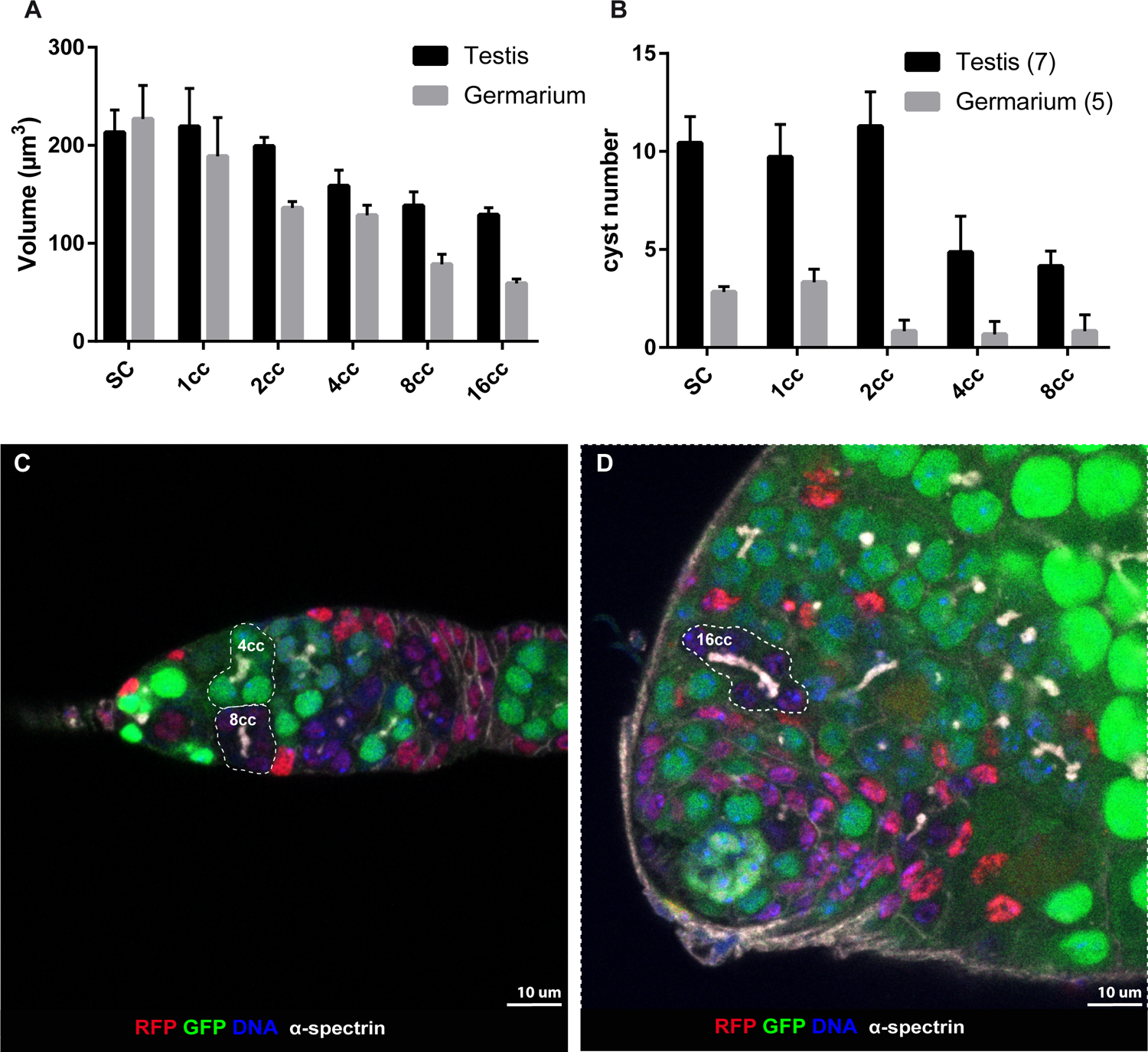
(A) Mean nuclear volume for each developmental stage obtained by live-imaging Nup::GFP/+; CID::RFP/+ germarium and testis. For each nucleus, the longest (D) and the smallest diameter (d) were determined by measuring the distance between two diametrically opposed Nup::GFP signals on projected images along the x-y axis. The height of the nucleus (h) was determined from z-series starting from the first Nup foci detected until the last one. Nuclear volume (μm3) was calculated using the formula: V= 4 x D x d x h x π/3. The number of analyzed nuclei (n) is indicated below each stage. Centre and error bars are mean ±SD. (B) Average number of cysts per stage in a testis (n = 7) and ovary (n = 15) SC: Stem Cell; 1 cc: Gonioblast (testis) or Cystoblast (Ovary); 2cc: 2 cell-cyst; 4cc: 4 cell-cyst; 8cc: 8 cell-cyst. (C,D) Selected projections from *yw^1118^ hs-FLP; FRT40A-GFP/ FRT40A-mRFP* used to generate mosaic germarium (C) and testis (D) stained for GFP (green), RFP (red), fusome (α–spectrin, white) and DNA (DAPI, blue). Having red, green or both color cysts, makes it easier to count the number of cysts at each stage in testis and ovaries. Scale bars are 10 µm.

## Supplementary Video Legends

**Supplementary Video 1.** Dynamics of centromere clusters in mitotic region of the testis. Time lapse microscopy (spinning disc) of a testis expressing the centromere marker CID::RFP (red) and the fusome marker MUD::GFP (green). One germinal stem cell (GSC) is identified by its position close to the hub (yellow circle). Two nuclei of an 8-cell cyst (8cc), whose cells are linked by a branched-shaped fusome, demonstrating that they are from the same cyst. Arrow points towards rotating centromeres cluster in an 8cc. Frames were taken every 10 seconds. The video is shown at 7 frames/s (MPEG4).

**Supplementary Video 2.** Centromere dynamics in living wild-type GSC. Time lapse microscopy (spinning disc) of a testis expressing the centromere marker CID::RFP (red). The movie is shown at 7 frames/s (MPEG4).

**Supplementary Video 3.** Centromere dynamics in living wild-type in living 8-cell cyst. Time lapse microscopy (spinning disc) of a testis expressing the centromere marker CID::RFP (red). The movie is shown at 7 frames/s (MPEG4)

**Supplementary Video 4.** *Dynein* loss of function in *Dhc64C ^3-2^/Dhc64C ^6-12^* mutant leads to inhibition of CID foci dynamics in living 8-cell cysts. Time lapse microscopy (spinning disc) of a *Dhc64C ^3-2^/Dhc64C ^6-12^* mutant testis expressing the centromere marker CID::RFP. Frames were taken every 10 seconds. The movie is shown at 7 frames/s (MPEG4).

**Supplementary Video 5.** Colcemid treatment leads to inhibition of CID foci dynamics in living 8-cell cysts. Time lapse microscopy (spinning disc) of a colcemid-treated testis expressing the centromere marker CID::RFP. Frames were taken every 10 seconds. The movie is shown at 7 frames/s (MPEG4).

**Supplementary Video 6.** CID-RFP and C(3)G-GFP remain in close proximity in living testis. Time lapse microscopy (spinning disc) of a testis expressing the centromere marker CID::RFP (red) and the C(3)G::GFP (green). Frames were taken every 30 seconds. The movie is shown at 7 frames/s (MPEG4).

## EXPERIMENTAL PROCEDURES

### Experimental Model and Subject Details Drosophila melanogaster

Flies were maintained on standard medium in 25°C incubators on a 12 h light/dark cycle. All *koi* rescue experiments were carried out at 29°C. Wild-type controls alone and in combination with additional transgenes of fluorescently tagged proteins were in a *w^1118^* background. The shRNA for white was used as control for knock-down experiments because *white* is not expressed during oogenesis and spermatogenesis.

### Fly stocks and genetics

For testing mutants, the following strains were used: *c(3)G^68^ (Page & Hawley, 2001), c(3)G^1^*(BDSC:606), *c(3)G^G5001^* (BDSC 30110), *cona^A12^/cona^f04903^(Page et al, 2008), koi^80^*((Kracklauer et al, 2007) BDSC:25105), *klar^mCD^*^4^ ((Technau & Roth, 2008) BDSC:25097), *Dhc64c^3−2^*/*Dhc64c^6−12^* (Gepner et al, 1996). The *c(3)G* deficiencies were: Df(3R)BSC569 (BDSC 25670), Df(3R)ED5705 (Bloomington 9152).

The shRNA lines were: for white, P{TRiP.GL00094}attP2 (BDSC: 35573); for CdK1/cdc2 P{TRiP.GL00262}attP2 (BDSC: 35350); for CycB P{TRiP.HMS01015}attP2 (BDSC: 34544), for twine P{TRiP.HMS00642} (Bloomington: 3044). *P{bam-GAL4:VP16}* (a gift from M. Fuller, Stanford University School of Medicine, CA, USA, and BDSC, #4937) was used as driver.

Live-imaging experiments were done using: CID::RFP (Schuh et al, 2007), Mud::GFP (Bosveld et al, 2016), protein trap Koi::GFP (y[1] w[*]; P{w[+mC]=PTT-GB}koi[CB04483] BDSC: 51525); *svv::GFP (Mathieu et al, 2013)* and *tmod::GFP* (Lighthouse et al, 2008); Nup107-GFP (Katsani et al, 2008).

### RNA extraction from ovaries and testis

For RT-qPCR, w^1118^ flies were used. Ovaries and testis were dissected and collected in cold PBS. Tissues were homogenized with a pestle and RNA was extracted following the standard protocol from the RNeasy Micro Kit (QIAGEN).

### RT–qPCR

cDNA was prepared from 0.5 µg total RNA following standard manufacturer’s protocol using random primers, 10 mM dNTP mix, 5X first strand buffer, 0.1 M DTT, 40 U RNaseOUT, 200 U SuperScript III RT (Invitrogen). Typically, 10% of the reaction was used as template. Real time PCR was carried out using SYBR Green Supermix (Bio-Rad) with a Roche LightCycler II 480. Cycle threshold (C _(T)_) values were determined by the software. Calculation of relative RNA levels were done using the 2[-ΔΔ C _(T)_] method (Livak & Schmittgen, 2001), where the C _(T)_ values were normalized to those of *rp49* in the same sample. Values used where the means of triplicate repeats.

Primers used in this study are:

*hth*: F 5’ AGCAAAGGCTTCCCAATACA 3’; R 5’ CCGTGTGGAGTCCTTAGATTTAC 3’

*bam:* F 5’ CTGCATATGATTGGTCTGCACGGC 3’; R

5’CCCAAATCGGCGGTCAGGTGATC 3’

*C3G1*:F 5’ GCACCGTTAGTGGATGCAG 3’; R 5’GGAGCCACTAGCGGAGTTCT 3’

*C3G3:* F 5’ TTTCCGGTTGCACCTTAGTAG 3’; R 5’

TAACTGCCTGATCGACCAAC 3’

*C3G4*: F 5’ GTGATCATGATACGCTAAAGAGC 3’; R 5’

ACTTTGGCCGTGGATAATTC 3’

*rp49*: F 5’ ATCTCGCCGCAGTAAACGC 3’; R 5’ CCGCTTCAAGGGACAGTATCTG 3’

### Immunohistochemistry

For confocal microscopy, testis and ovaries were dissected in PBS, fixed in 4% PFA–PBS, and then permeabilized in PBT (0,2% Triton) for 30 min. Samples were incubated overnight with primary antibodies in PBT at 4 °C, washed 4 × 30 min in PBT, incubated with secondary antibody for 2 h at room temperature, washed 4 × 30 min in PBT. DAPI (1:500) was added during the last wash and then mounted in CityFluor.

For STED and Airyscan microscopy, testes were dissected in PBS, fixed in 4% PFA–PBS, permeabilized in PBT (0.2% Triton) for 30 min, then in PBT (0,4% Triton) + 10% BSA) for 1 hour. Samples were incubated for 72 hours with primary antibodies at 4 °C, washed 4 × 30 min in PBT, incubated with secondary antibody for 2 h at room temperature, washed 4 × 30 min in PBT. DAPI (1:500) was added during the last wash and then mounted in CityFluor. For FISH experiments, testes were dissected in PBS, fixed in 4% PFA in 1X fix buffer (100 mm potassium cacodylate, 100 mm sucrose, 40 mm sodium acetate, and 10 mm EGTA).

Testes were then rinsed three times in 2X SSCT and incubated with the AACAC and dodeca probes which target the pericentromeric regions of the 2^nd^ and 3rd chromosome, respectively, as previously described (Christophorou et al, 2013). Testes were then rinsed in 2X SSCT, twice in PBST and process for immunostaining as described above for confocal microscopy. The following primary antibodies were used: mouse anti-C(3)G 1A8-1G2 (1:500) (gift from S. Hawley, Stowers Institute, USA), rat anti-Cid (1:1,000) (gift from C. E. Sunkel, Universidade do Porto, Portugal), rabbit anti-α-Spectrin (1:1,000 and 1:500 when used with FISH) (gift from R. Dubreuil, University of Chicago, USA), rat anti-Klaroid (1:200), guinea pig anti-Klarsicht (1:200) (gifts from M. Welte, University of Rochester, USA and J. Fischer, University of Texas, USA). Secondary antibodies conjugated with Cy3, Cy5, FITC (Jackson laboratories) and STAR RED (abberior) were all used at 1:200.

### Colcemid treatments

Colcemid was prepared at a concentration of 0.2 mg ml^−1^ in 1% saccharose, mixed with dry yeast and then added to the fly medium. To assay pairing and clustering of centromeres in fixed samples, long-term drug treatments lasted 48 h. Freshly prepared drug was added every 12 h. For live imaging, adult flies were fed for four hours with colcemid-containing media. Live imaging was performed as described below.

### Image acquisition

Live and fixed imaging of testes and ovaries was done from 3-5 day-old flies For fast acquisitions of centromeres, all dissections were done in oil (10S, Voltalef, VWR). The muscular sheath around each ovariole was removed and germaria were made to stick to coverslips. Live-imaging was done using an inverted spinning-disc confocal microscope (Roper/Nikon) operated by Metamorph software on an inverted Nikon Eclipse Ti microscope coupled to an Evolve EMCCD camera (Photometrics). All images were acquired with the PlanApo 60×/1.4 NA Oil objective with ×1.5 auxiliary magnification. Single-position videos in testis and germarium were acquired for 8 min at 25 ± 1 °C, with a 10 s temporal resolution (11-slice *Z*-stack, 0.5 μm per slice).

For long cell cycle live imaging of females versus males, ovaries and testis were dissected in Schneider medium (Sigma-Aldrich) supplemented with 15% fetal calf serum and 0.6% penicillin-streptomycin (Invitrogen). Muscle sheath from germaria was manually removed. Several *w^1118^; svv::GFP/+; tmod::GFP/+* testis and ovarioles were each placed in a small drop of Growth Factor Reduced Matrigel (Corning) on a glass coverslip culture dish (P35G-1.0-14-C MatTeck), filled with the medium, covered with a gas permeable membrane, and sealed with oil. Imaging was performed using a DeltaVision (Applied Precision) microscope system equipped with an Olympus 1670 inverted microscope and a CoolSNAP HQ CCD 40×1.42 camera. Confocal images (55 sections of 1.3 μm per time point) were collected every 10 minutes. Tracking of cells was done manually using Fiji (ImageJ) processing program.

For cell cycle rescue live imaging, a protocol adapted from (Burnett et al, 2018b) was used. *w; svvGFP; bam>shRNA* ovaries were dissected in supplemented Schneider medium, and transferred onto a round 25 mm coverslip previously coated with 3-(trimethoxysilyl) propyl methacrylate (Sigma-Aldrich). Medium was then replaced with 10% PEG-DA hydrogel solution (esibio) containing 0.1% I2959 (photoinitiator, Sigma-Aldrich). Another coverslip treated with water repellent (Rain-X) was placed over the hydrogel droplet. The coverslip/coverslip sandwich was then exposed to a UV Transilluminator for 30 s (312 nm) for gelation. The upper coverslip was then removed and the lower coverslip supporting the hydrogel disc placed into a Chamlide chamber filled with medium (Burnett et al, 2018a).

Videos were collected with an inverted spinning-disc confocal microscope (Roper/Nikon) operated by Metamorph on an inverted Nikon Eclipse Ti microscope coupled to an Evolve EMCCD camera (Photometrics). All images were acquired with a PlanApo 60×/1.4 NA oil objective with ×1.5 auxiliary magnification. Single-position videos were acquired for 8 min, with a 10 s temporal resolution (11-slice *Z*-stack, 0.5 μm per slice). All images were acquired at 29°C.

Confocal images of fixed germaria and testes were obtained with a Zeiss LSM 980 NLO confocal. All images were acquired with a PlanApo 63×/1.40 NA oil objective at 0.5 μm intervals along the *z*-axis operated by ZEN 2012 software.

Super-resolution STED imaging was performed with a STEDyCON (Abberior Instruments) mounted at the camera port of an AxioObserver (Zeiss) microscope equipped with a 100 × objective (APO 100x/1.4 NA Oil). Star Red was imaged with excitation wavelength of 640 nm and time-gated fluorescence detection between 650 and 720 nm. The STED laser wavelength of 775 nm and a pulse width of ∼ 1–7 ns were used. The pinhole was set to 1.13 AU. The images were further processed (black/white inversion) with the Fiji software 103.

### Data analysis

For quantification of CID foci on fixed tissue, we counted the number of distinguishable CID foci within each single nucleus. For quantification in live tissue, we consider all CID foci at a given (t) projection where the number of CID foci was maximal. In all figures, micrographs represent the projections of selected z-series taken from the first CID foci signal until the last one. For FISH experiments, the 3D distances between the AACAC foci and between the dodeca foci were measured as described (Christophorou et al, 2013). Pericentromeric regions of chromosomes were considered as paired when only one foci was visible or when two foci were separated by a distance less than 0.70 µm, and as unpaired when ≥ 0.70 µm. To assign the number of CID dots to a cyst developmental stage, we used α-Spectrin immunolabelling in fixed tissue and live MUD::GFP expression.

Three-dimensional tracking of CID foci was performed using Imaris software (Bitplane). The CID::RFP signal was tracked using the ‘spots’ function with an expected diameter of 0.6 μm. Automatically generated tracks were edited manually to eliminate inappropriate connections (*e.g.* connections between foci in different nuclei or between foci of different sizes or intensity, multiple assignments or multiple spots assigned to the same focus). To remove global movements of the germarium, each nucleus containing a CID::RFP focus was assigned to the nearest fusome. Then, the position of the reference fusome was subtracted from each CID::RFP focus for each time point to obtain the relative tracks. These relative tracks were then compiled using a custom MATLAB (MathWorks) routine that computes the minimum volume of the ellipsoid that encloses all of the three-dimensional points of the trajectory.

### Analysis of centromere trajectories

Positions of individual centromeres were tracked every 10 s during 8 min to quantify the volume covered by each centromere. This raw volume was then corrected both for overall movements of the tissue and for variations in total nuclear volume. First, we subtracted the motion of the germarium using the position of the fusome as a reference within each cyst. Second, to take into account the significant decrease of the nuclear volume from GSCs to 16-cell cysts (see Supplementary Fig.3A-B), we computed the relative volume, which is the raw volume divided by the mean value of the nuclear volume at each stage. Finally, we normalized the durations of each track by calculating the relative covered volume per second.

### Mean Cyst number estimation

To generate *yw^1118^ hs-FLP; FRT40A-GFP/ FRT40A-mRFP* flies, we crossed *yw^1118^ hs-FLP; FRT40A-GFP* with *yw hs-FLP; FRT40A-mRFP.* The resulting larvae were subjected to 1 hour heat shocks for 3 days to activate the flipase. This allows recombination at the FRT sites, to generate clones of red (homozygous mRFP), green (homozygous GFP) or green and red (mRFP / GFP) colors. Mosaic adult ovaries and testes having red, green or green and red cells, makes it easier to count the number of cysts at different stages. We used the mean cyst number to calculate cell cycle length.

### Cell cycle length estimation

To determine the cell cycle of germ cells in males and females we used *w^1118^; svv::GFP; tmod::GFP* flies. Here *tmod::GFP* served as a fusome marker and *svv::GFP,* which associates with centromeres during mitosis, allowed counting the number of mitosis. For example, 1 testis with ± 10.4 GSCs, we calculated the total amount of time GSCs were imaged (10.4 GSC x 15.8 hours) and divided it by the total number of GSC divisions observed during that time (8 cell divisions) to obtain a mean cell cycle duration for GSCs of 20.6 hours (10.4 x 15.8/8). We calculated similarly the mean cell cycle time for each stage (Figure 6A-B) (Sheng & Matunis, 2011). The length of each movie was determined by the last germline mitosis observed at any stage.

To measure the effect of shRNA lines on cell cycle length, *svv::GFP/CyO; bamGal4/TM3Ser* females were crossed shRNA males. The resulting progenies of svv::GFP; bam>shRNA females were dissected in hydrogel, as described above. Live-images were analyzed as above.

### Reproducibility of experiments

Images in all Figures and Supplementary Figures are representative images of at least 3 independent experiments.

